# Surpassing the physical Nyquist limit to produce super-resolution cryo-EM reconstructions

**DOI:** 10.1101/675397

**Authors:** J. Ryan Feathers, Katherine A. Spoth, J. Christopher Fromme

**Affiliations:** Department of Molecular Biology and Genetics, Weill Institute for Cell and Molecular Biology, Cornell University, Ithaca, NY 14853 USA; Cornell Center for Materials Research, Cornell University, Ithaca, NY 14853 USA

## Abstract

The resolution of cryo-EM reconstructions is fundamentally limited by the Nyquist frequency, which is half the sampling frequency of the detector and depends upon the magnification used. In principle, super-resolution imaging should enable reconstructions to surpass the physical Nyquist limit by increasing sampling frequency, yet there are no reports of reconstructions that do so. Here we report the use of super-resolution imaging with the K3 direct electron detector to produce super-resolution single-particle cryo-EM reconstructions significantly surpassing the physical Nyquist limit. We also present a comparative analysis of a sample imaged at four different magnifications. This analysis demonstrates that lower magnifications can be beneficial, despite the loss of higher resolution signal, due to the increased particle numbers imaged. To highlight the potential utility of lower magnification data collection, we produced a 3.5 Å reconstruction of jack bean urease with particles from a single micrograph.

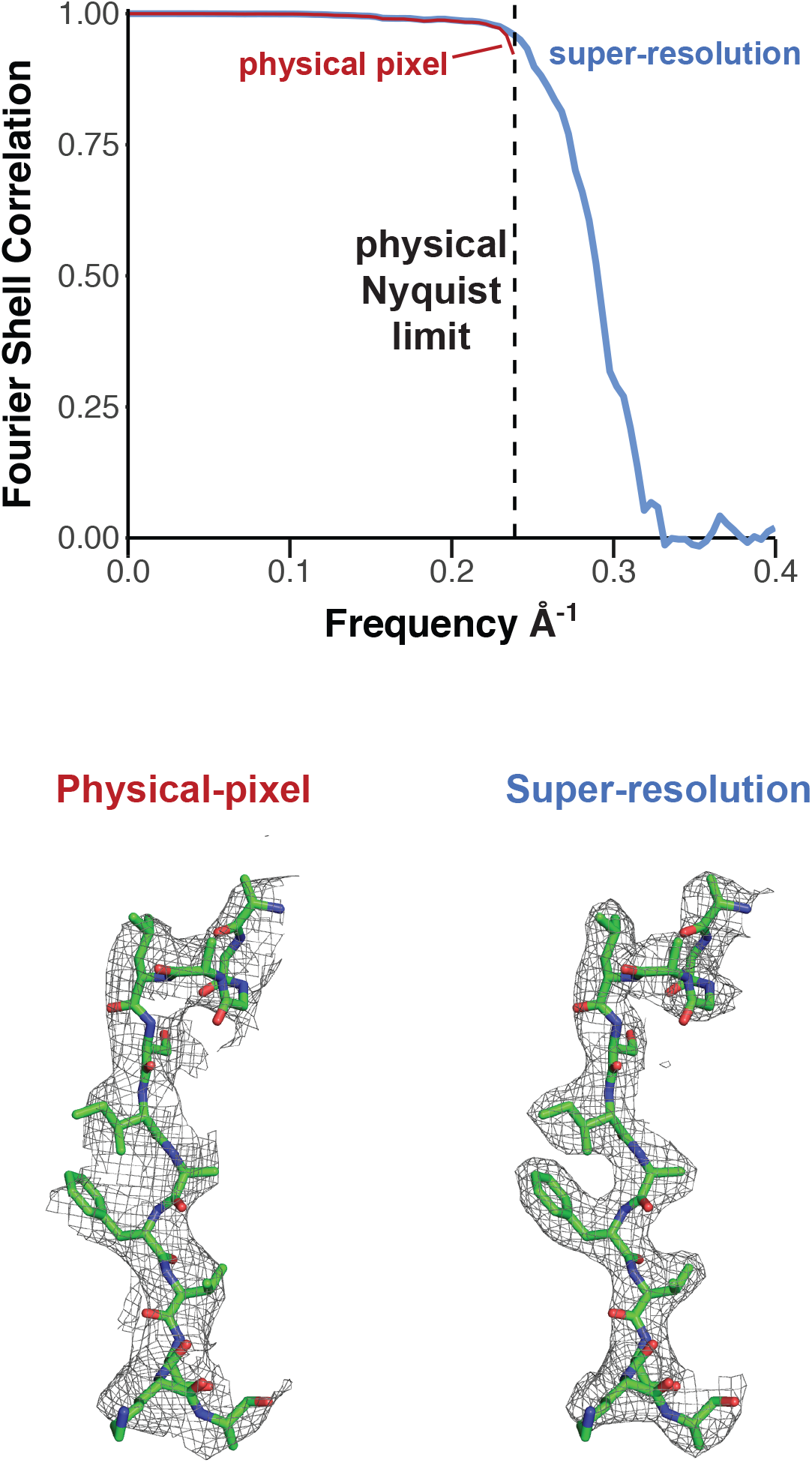

## Introduction

The use of single-particle cryo-electron microscopy (cryo-EM) as a structural biology technique has gained wide adoption in recent years, in large part due to the development of direct electron detectors and advances in computational methods (Cheng et al., 2015; Cheng, 2015; Nogales and Scheres, 2015; Cheng et al., 2016). Dozens of institutions worldwide have acquired the sophisticated EM tools required to collect high-resolution data for determining the atomic structures of macromolecules using single-particle analysis techniques. Due to the high cost of acquisition and maintenance of these tools, as well as high demand for their use, there are strong incentives to maximize the amount of data collected per unit time.

A typical 24-hour experiment on a high-resolution EM instrument involves ~4-8 hours of setup time and ~16-20 hours of data collection time, resulting in hundreds or thousands of image exposures. A key decision made by the experimenter is the magnification used for imaging. The magnification needs to be large enough to capture high-resolution information. However, the higher the magnification used, the smaller the field of view and therefore the fewer particle images that are captured per micrograph. As a consequence, using higher magnifications results in fewer particle images collected in the same time period.

The resolution of cryo-EM reconstructions is fundamentally limited to the Nyquist frequency, which is half the sampling frequency of the detector. For example, if data is collected using a magnification which produces an image on the detector such that each 1.5 Å of the sample spans one pixel of the detector (image pixel spacing or “pixel size” = 1.5 Å), then the Nyquist frequency is 1/3 Å^−1^ and the maximum achievable resolution of the reconstruction (“Nyquist limit”) is 3.0 Å. The achievable resolution is further constrained by aliasing. Aliasing arises when an image projected onto a detector contains information at higher resolution (frequencies) than the Nyquist limit. This leads to artifacts in the image that can limit the achievable resolution.

The advent of direct electron detectors for cryo-EM experiments and advances in data processing have resulted in a “revolution” of hundreds of new high-resolution macromolecular structures. These detectors can count individual electrons, greatly improving signal-to-noise (Bammes et al., 2012; Li et al., 2013b; McMullan et al., 2014; Ruskin et al., 2013; Cheng et al., 2016; Mendez et al., 2019). Fast readout rates enable exposures to be collected as movies which can be corrected for image blurring caused by sample motion. This motion correction results in a significant improvement in the information content at high resolution (Campbell et al., 2012; Li et al., 2013a).

Some direct electron detectors, such as the Gatan K2 and K3, can also generate “super-resolution” images. The super-resolution method takes advantage of the fact that a single electron is detected simultaneously by multiple adjacent pixels. The centroid of the electron can therefore be attributed to one quadrant of a physical pixel, resulting in a four-fold increase in the number of image pixels compared to physical detector pixels (Booth, 2012; Li et al., 2013b). At a given magnification, super-resolution data acquisition should reduce the effects of aliasing by effectively doubling the spatial sampling frequency.

Another limit to the high-resolution information available in an exposure is the detective quantum efficiency (DQE) of the detector, which is essentially a measure of the detector’s signal-to-noise performance. Although the precise shape of the DQE curve varies among different detectors, DQE declines as the spatial frequency approaches the Nyquist limit (McMullan et al., 2014, 2009; Milazzo et al., 2010; Ruskin et al., 2013). Using higher magnification ensures that the DQE is sufficiently high to capture high-resolution information. For super-resolution detectors, DQE is relatively low but non-zero in the super-resolution range. The general view of many in the field is that the DQE in the super-resolution range is low enough that it is not worth the added storage and computational costs associated with preserving images in super-resolution format; instead super-resolution images should be Fourier-cropped (“binned”). This reduces the negative effects of aliasing but discards the potentially useful super-resolution information.

In principle, super-resolution imaging should enable the resolution of cryo-EM reconstructions to surpass the physical Nyquist limit, and to approach what we term the “super-Nyquist” limit defined as half the sampling frequency of the super-resolution image. Indeed, super-resolution information was detectible in images collected on 2D protein crystals (Chiu et al., 2015), and therefore it was expected that super-resolution reconstructions should be possible. However, no 3D reconstructions with resolutions surpassing the physical Nyquist limit have been reported in the literature. Therefore, it remains unresolved whether single-particle reconstructions can be easily produced with resolutions that surpass the physical Nyquist frequency.

Here we report the production of super-resolution single-particle cryo-EM reconstructions that surpass the physical Nyquist limit by as much as 1.4-fold. With a physical pixel size of 1.66 Å we obtained a 2.77 Å reconstruction and with a physical pixel size of 2.10 Å we obtained a 3.06 Å reconstruction. We collected datasets at multiple magnifications to compare the tradeoffs between increasing particle numbers and decreasing high resolution signal. As expected, at the lowest magnification the resulting reconstructions appeared to be limited by low detector DQE at high frequencies. However, we identified a magnification range where lower magnifications produced reconstructions with the same resolution as those produced at higher magnifications, but required fewer exposures. As a proof of principle, we also used this approach to generate sub-4 Å reconstructions of jack bean urease using single exposures.

## Results

Typical high-resolution single-particle cryo-EM experiments utilize physical pixel sizes of approximately 1.0 Å. For a medium-sized sample with dense particle distribution, this might result in ~400 particles per micrograph, and 500 exposures will result in a dataset with ~200,000 particles (Table 1). Samples are typically imaged after being applied to grids coated with a foil substrate consisting of an array of ~1-2 μm holes, with a single exposure taken per hole (alternative data collection strategies are addressed in the Discussion). The result is that most of the particles present in a hole are not actually imaged. Using a lower magnification results in more particles per exposure and therefore per dataset (Figure S1A and Table 1). However, the accompanying increase in the pixel size of the image results in a corresponding shift of the DQE, consequently degrading signal at high spatial frequencies (Figure S1B). Researchers therefore balance the need for high resolution information with the need for large numbers of particles when choosing a magnification for imaging.

**Table 1:** Estimated number of particles per micrograph at different magnifications.

| Nominal Magnification | Detector | Image Physical Pixel Size | Physical Nyquist | Super-Nyquist | Mag detector area | ~particles / micrograph |
| --- | --- | --- | --- | --- | --- | --- |
| 79kX | K2 | 1.03 Å | 2.06 Å | 1.03 Å | 15.1 million Å<sup>2</sup> | 400 |
| 63kX | K2 | 1.29 Å | 2.58 Å | 1.29 Å | 23.7 million Å<sup>2</sup> | 630 |
| 100kX | K3 | 0.83 Å | 1.66 Å | 0.83 Å | 19.6 million Å<sup>2</sup> | 520 |
| 79kX | K3 | 1.03 Å | 2.06 Å | 1.03 Å | 25.0 million Å<sup>2</sup> | 660 |
| 63kX | K3 | 1.29 Å | 2.58 Å | 1.29 Å | 39.2 million Å<sup>2</sup> | 1000 |
| 49kX | K3 | 1.66 Å | 3.32 Å | 1.66 Å | 64.9 million Å<sup>2</sup> | 1700 |
| 39kX | K3 | 2.10 Å | 4.20 Å | 2.10 Å | 103.9 million Å<sup>2</sup> | 2750 |

The “~particles / micrograph” numbers in this table assume a uniform sample distribution resulting in 400 particles per K2 image at 1.03 Å/pixel
The K2 detector is a 3838 x 3710 array of pixels
The K3 detector is a 5760 x 4092 array of pixels

### Production of super-resolution single-particle 3D reconstructions

We began with the goal of exploring whether super-resolution imaging could be used to produce single-particle 3D reconstructions with 0.143-FSC resolutions that exceed the physical Nyquist limit. We therefore chose a magnification at which the resolution of a reconstruction using physical pixels would likely be constrained by the physical Nyquist limit. We imaged jack bean urease frozen in vitreous ice using a Talos Arctica operating at 200 kV with a Gatan K3 mounted on an imaging filter and a nominal magnification of 49,000x, resulting in a physical pixel size of 1.66 Å. Although the structure of this protein has been determined by X-ray crystallography (Balasubramanian and Ponnuraj, 2010), no EM reconstruction has been reported. Due to the lower magnification and fairly dense particle distribution, each micrograph contained ~2,200 particles. (Figure 1A).

**Figure 1:**
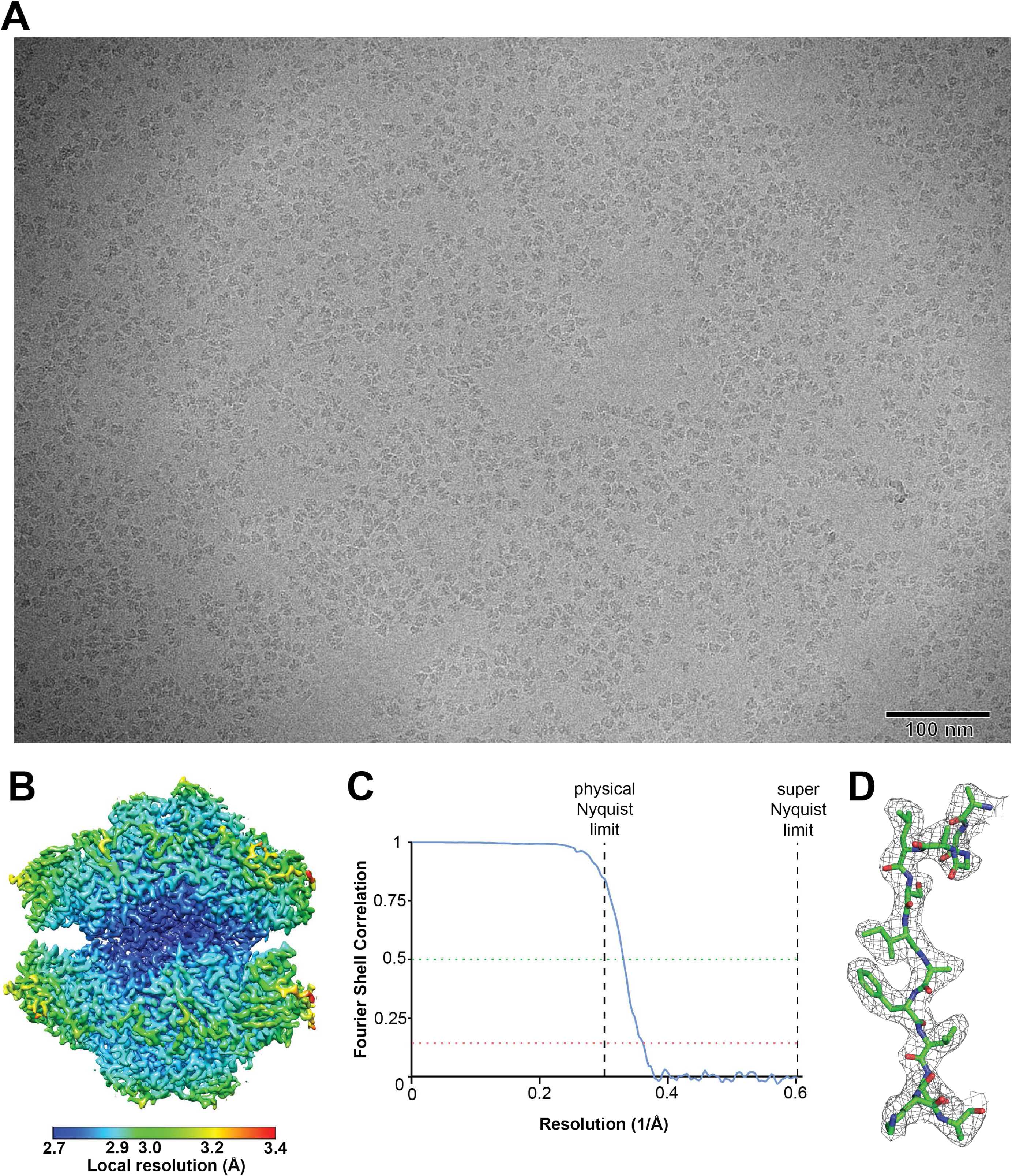
2.77 Å super-resolution reconstruction of jack bean urease surpasses the physical Nyquist limit of 3.32 Å. A) Example motion-corrected micrograph imaged at 49kX nominal magnification resulting in a physical pixel size of 1.66 Å and physical Nyquist limit of 3.32 Å. The width of the magnified image is 956 nm. The image has been low-pass filtered to 20 Å. B) Final sharpened reconstruction colored by local resolution. C) FSC curve for the final reconstruction particle set. The 0.143 and 0.5 FSC cutoffs are shown as dotted lines, the physical Nyquist and super-Nyquist frequencies are shown as dashed lines. D) Example electron density of the final sharpened 3D reconstruction overlaid on the atomic model. See Figure S2 for a graphical depiction of the data processing steps.

We used Serial-EM software (Mastronarde, 2005; Schorb et al., 2019) to automatically collect 240 super-resolution exposures. We did not bin the super-resolution micrographs during data collection or motion-correction, therefore the image pixel size of the micrographs was 0.83 Å. Relion 3.0 (Zivanov et al., 2018) was used for all data processing steps, including wrappers to MotionCor2 (Zheng et al., 2017) for motion correction and dose-weighting of movies and to GCTF (Zhang, 2016) for initial per-micrograph defocus estimation. To generate a 3D reference for template-based autopicking, we first performed 3D refinement with a small subset of the data, using a 60 Å low-pass filtered crystal structure of urease (Balasubramanian and Ponnuraj, 2010) as an initial reference model. Autopicking using this 3D template yielded ~359,000 particles from the entire dataset. Particle images were first extracted using a Fourier-cropped pixel size of 1.66 Å, equivalent to the physical pixel size, and 3D-refinement of the entire particle set resulted in a 3D reconstruction with a masked 0.143 Fourier-shell correlation (FSC) resolution (Rosenthal and Henderson, 2003) that reached the physical Nyquist limit of 3.32 Å without any additional processing (see Figure S2 for FSC curve).

Particle images were then re-extracted using the super-resolution pixel size of 0.83 Å. Several rounds of 3D refinement interspersed with per-particle CTF refinement, Bayesian particle polishing, beam-tilt estimation (Zivanov et al., 2018, 2019), and 3D-classification produced a final subset of ~56,000 particles generating a 3D reconstruction with a masked 0.143 FSC resolution of 2.77 Å (Figures 1B,C, S2, and Table 2). We note that using a 0.5 FSC cutoff, the resolution was 3.0 Å, and the electron density map showed clear side-chain densities consistent with a ~2.8 Å reconstruction (Figure 1D). To assess the suitability of the reconstruction for atomic model refinement we manually modified the X-ray crystallographic atomic model of urease to fit the sharpened map and performed real-space refinement (Afonine et al., 2018b), producing a refined atomic model with excellent geometry and a model-map 0.5 FSC of 2.9 Å (Figure 1D and Table 3). The sharpened map was of sufficient quality to enable *de novo* atomic model building. In fact, virtually all of the protein backbone and more than half of the sidechains were automatically modeled correctly using the automated “map-to-model” procedure (Terwilliger et al., 2018) within the Phenix software package (Adams et al., 2010) (Table 3).

**Table 2:**
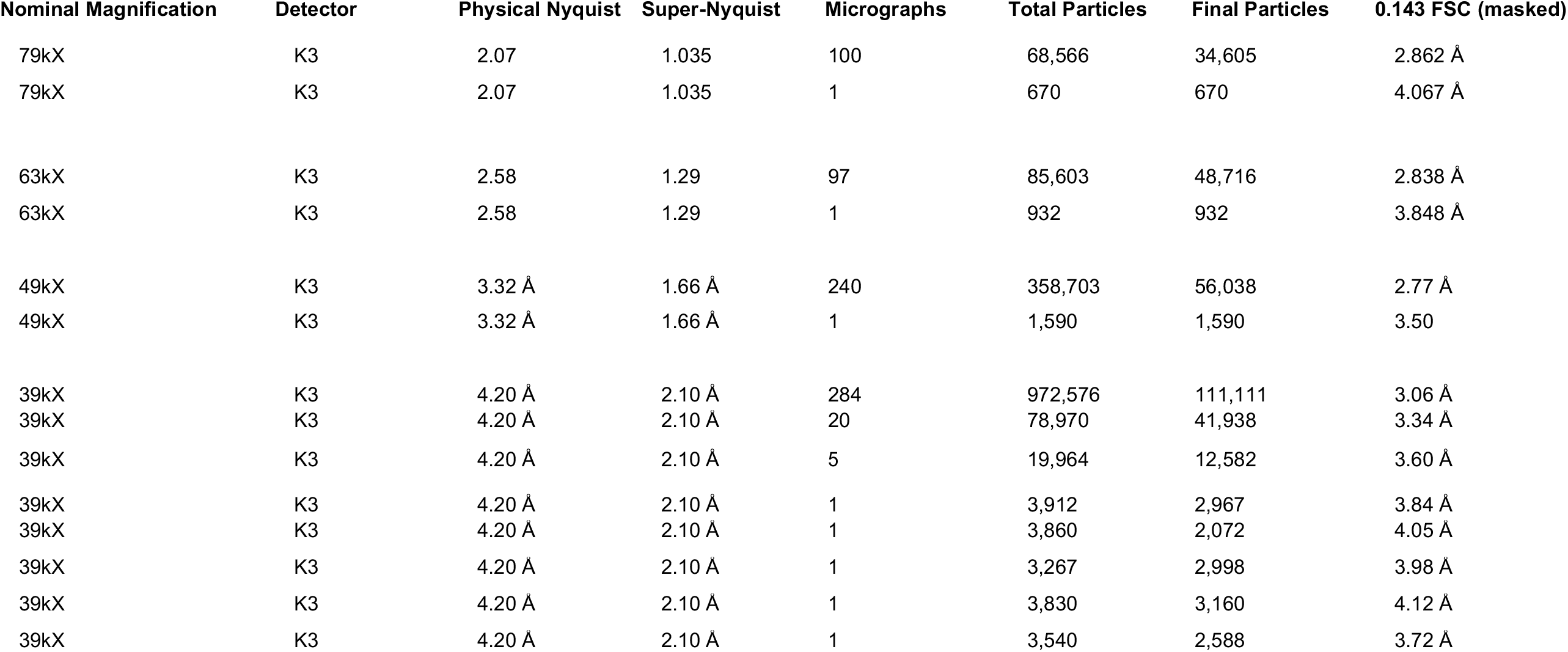
Datasets and reconstructions.

**Table 3:** Lower magnification atomic model statistics.

|  |  |  |  |
| --- | --- | --- | --- |
| Imaging conditions |  |  |  |
| Nominal magnification: | 49kX | 39kX | 39kX |
| Physical pixel size: | 1.66 Å | 2.10 Å | 2.10 Å |
| Super-res pixel size: | 0.83 Å | 1.05 Å | 1.05 Å |
| Micrographs in dataset: | 240 | 260 | 1 |
| Particles used for final model: | 56,038 | 111,111 | 2,967 |
| 0.143 FSC (masked): | 2.77 Å | 3.06 Å | 3.84 Å |
| Model-map 0.5 FSC: | 2.8 Å | 3.1 Å | 3.8 Å |
| Model-map masked CC | 0.89 | 0.84 | 0.83 |
| Ramachandran plot (%) |  |  |  |
| Favored | 97.05 | 97.17 | 92.97 |
| Allowed | 2.95 | 2.83 | 7.03 |
| Outliers | 0.00 | 0.00 | 0.04 |
| Clash score | 3.55 | 5.00 | 8.45 |
| Bonds (RMSD) |  |  |  |
| Length (Å) | 0.004 | 0.004 | 0.005 |
| Angles (°) | 1.174 | 0.593 | 0.699 |
| Rotamer outliers (%) | 0.00 | 0.00 | 0.00 |
| MolProbity score | 1.31 | 1.41 | 1.91 |
| Map-to-model automated building |  |  |  |
| Close (%) | 95.1 | 93.8 | 50.9 |
| Matching sequence (%) | 55.2 | 44.9 | 8.0 |

To determine if even lower magnifications could be used to obtain high-resolution reconstructions, we collected another dataset on the same sample at a nominal magnification of 39,000x, with a physical pixel size of 2.10 Å (Figure 2A). We collected 284 super-resolution movie exposures and autopicked ~973,000 particles, with most images containing ~3500-3900 particles. 2D-classification (Figure 2B) was used to remove ~128,000 “junk” particles. The remainder of the data processing procedure was similar to that described above (Figure S3), resulting in a reconstruction with a masked resolution of 3.06 Å at the 0.143 FSC cutoff (3.30 Å at 0.5 FSC cutoff) using ~110,000 particles (Figure 2C,D and Table 2). Again, the electron density map quality was consistent with this resolution value (Figure 2E) and suitable for *de novo* model building (Table 3), confirming that the physical Nyquist limit of 4.20 Å was significantly surpassed.

**Figure 2:**
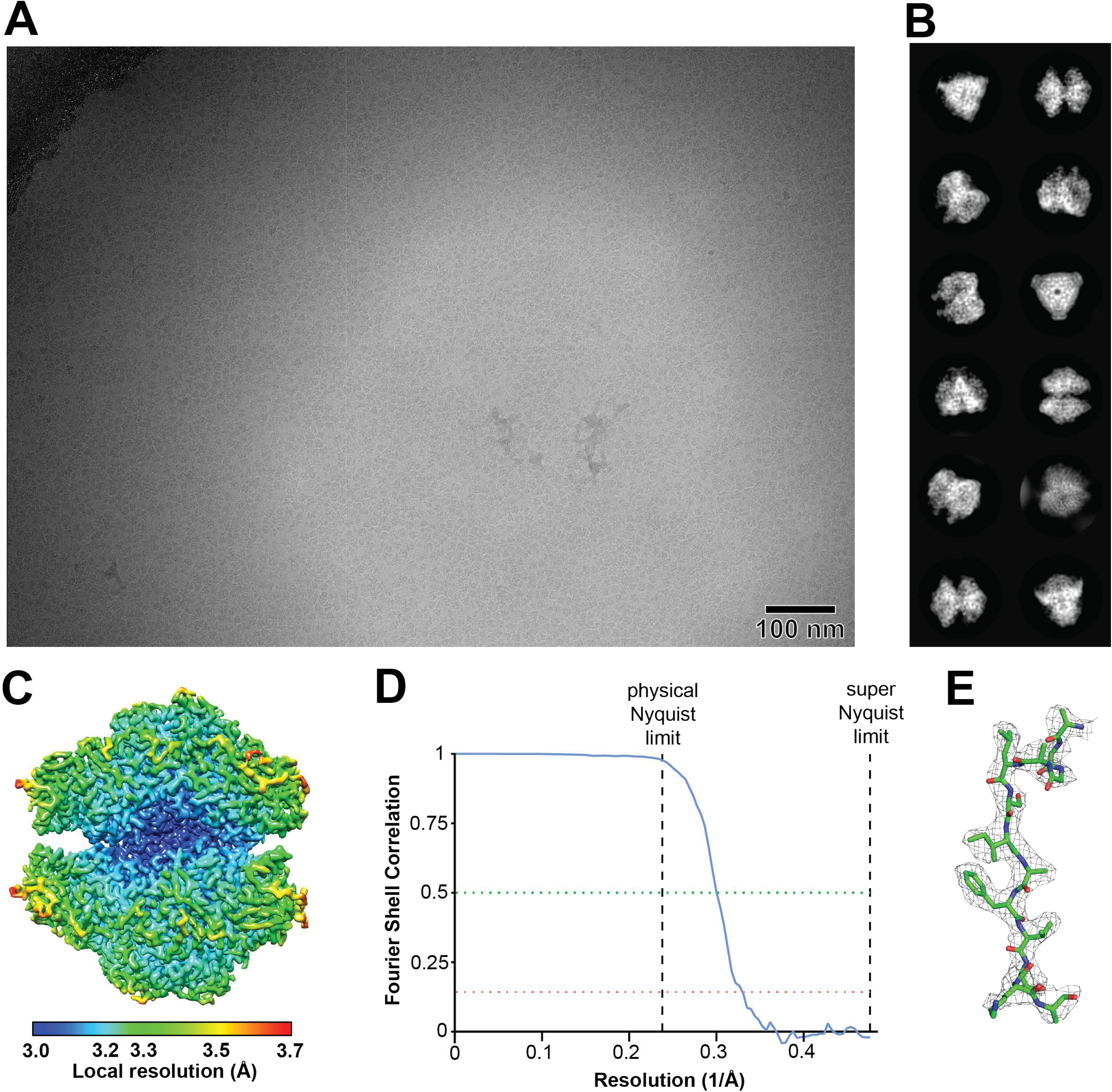
3.06 Å super-resolution reconstruction of jack bean urease surpasses the physical Nyquist limit of 4.20 Å. A) Example motion-corrected micrograph imaged at 39kX nominal magnification resulting in a physical pixel size of 2.10 Å and physical Nyquist limit of 4.20 Å. The width of the magnified image is 1,210 nm. The vertical white line is an unexplained imaging artefact. The gradient of ice thickness within the hole is visible as a contrast gradient. The edge of the hole is visible in the upper left corner of the image. The image has been low-pass filtered to 20 Å. B) Selected classes after initial 2D-classification to remove junk particles. Individual box edge width is ~230 Å. C) Final sharpened 3D-reconstruction colored by local resolution. D) FSC curve for the final reconstruction particle set. The 0.143 and 0.5 FSC cutoffs are shown as dotted lines, the physical Nyquist and super-Nyquist frequencies are shown as dashed lines. E) Example electron density of the final sharpened 3D reconstruction overlaid on the atomic structure. See Figure S3 for a graphical depiction of the data processing steps.

We considered the fact that ~110,000 is a rather large number of particles for reconstruction of a molecule with D3 (6-fold) symmetry. Additional 3D-classification to obtain a smaller subset of particles resulted in a 3.19 Å reconstruction using ~47,000 particles (data not shown). For comparison, the dataset imaged at 49 kX described above produced a 2.8 Å resolution reconstruction with ~56,000 particles. Taken together, these results suggest that these ~3.1 - 3.2 Å reconstructions using a 2.1 Å physical pixel size have reached the practical information limit, likely constrained by insufficient DQE at higher frequencies. We therefore conclude that super-resolution imaging with the K3 can be used to obtain reconstructions with 0.143-FSC resolutions reaching ~4/3 the physical Nyquist frequency limit.

### Super-resolution information significantly improves a physical Nyquist frequency-limited reconstruction

To directly assess the extent to which super-resolution information contributed to the reconstruction, we generated two 3D-reconstructions from the same set of particles, using the same refined particle orientations and weights, either with or without the super-resolution information intact (Figure 3A). We used with the final set of ~110,000 particles used to produce the 39kX reconstruction; this low magnification was chosen to maximize the potential effect of the super-resolution information. We re-extracted these particles and Fourier-cropped the particle images to the physical pixel size of 2.1 Å. The resulting particle images do not contain super-resolution information, other than the beneficial effect of reducing aliasing artifacts. These particles were refined and used to generate a 3D-reconstruction whose resolution is clearly Nyquist limited to 4.2 Å (Fig 3B). The particles were then re-extracted without Fourier-cropping, therefore preserving the super-resolution information, and a 3D-reconstruction was produced without any additional refinement. This reconstruction had a 0.143-FSC resolution of 3.2 Å (Fig 3B), slightly worse than the 3.1 Å resolution produced above, likely for two reasons. First, it was necessary to avoid Bayesian polishing during this process in order to directly compare the same refined particles with and without Fourier-cropping, and second, the super-resolution information was not used during the refinement procedure.

**Figure 3:**
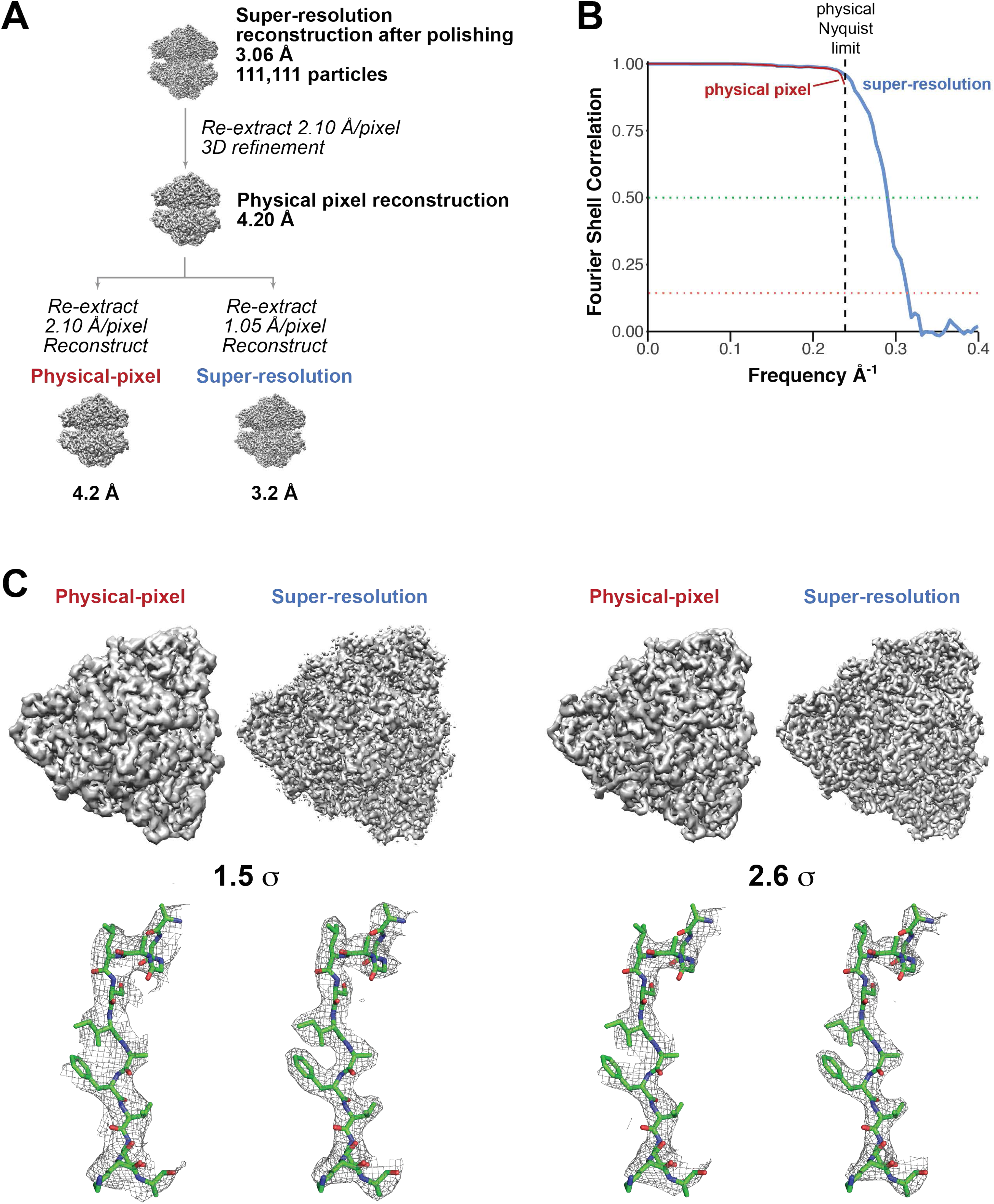
Direct comparison of reconstructions with and without super-resolution information. A) Flowchart indicating how reconstructions were produced from the same particles with and without super-resolution information. B) FSC curves for the resulting reconstructions. C) Comparison of the resulting cryo-EM reconstruction density maps, contoured at 1.5 σ (left) and 2.6 σ (right).

Direct comparison of the reconstructions produced with and without the super-resolution information revealed marked differences in the level of detail observable in the cryo-EM density (Figure 3C). Sidechain density was much more clearly resolved in the reconstruction produced with super-resolution information. This provides additional evidence indicating that useful information was obtained at super-resolution frequencies.

### Production of 3D-reconstructions from single micrographs

Given the very large numbers of particles obtained per image at this magnification, we were curious to determine the smallest number of exposures that were required to produce a useful 3D-reconstruction. Using the same 39,000x magnification dataset, we reprocessed the data using smaller numbers of images. Importantly, we did not select the best particle subsets from the refined larger dataset, or select the best subset of micrographs. Instead we simply used the first exposures we collected and repeated the data processing procedure beginning with particle autopicking. We processed datasets with sizes of 20 and 5 micrographs. Surprisingly, both of these datasets resulted in 3D-reconstructions with masked 0.143 FSC resolutions, 3.34 Å and 3.60 Å, surpassing the physical Nyquist limit (Table 2).

We then chose two individual micrographs from the 39kX dataset, selecting from among the first twenty those with the lowest estimated defocus values (0.8 and 0.9 μm) that also displayed good contrast. We processed each of these single micrographs as an independent dataset. Particles from one micrograph (0.8 μm defocus) generated a 3D-reconstruction with a masked 0.143 FSC resolution of 4.12 Å, while particles from the other micrograph (0.9 μm defocus, shown in Figure 2A) generated a 3D-reconstruction with a masked 0.143 FSC resolution of 3.84 Å (Figures 4A,B,C, S4A, and Tables 2 and 3). We then selected three more individual micrographs from the larger 39,000x magnification dataset to process as independent datasets, and each of these produced 3D-reconstructions with similar masked 0.143 FSC resolutions: 4.05 Å, 3.98 Å, and 3.72 Å (Figure 4B and Table 2). Remarkably, these five individual micrographs each resulted in a 3D-reconstruction that surpassed the physical Nyquist limit of 4.20 Å, although only three out of the five did so convincingly (Figure 4B).

**Figure 4:**
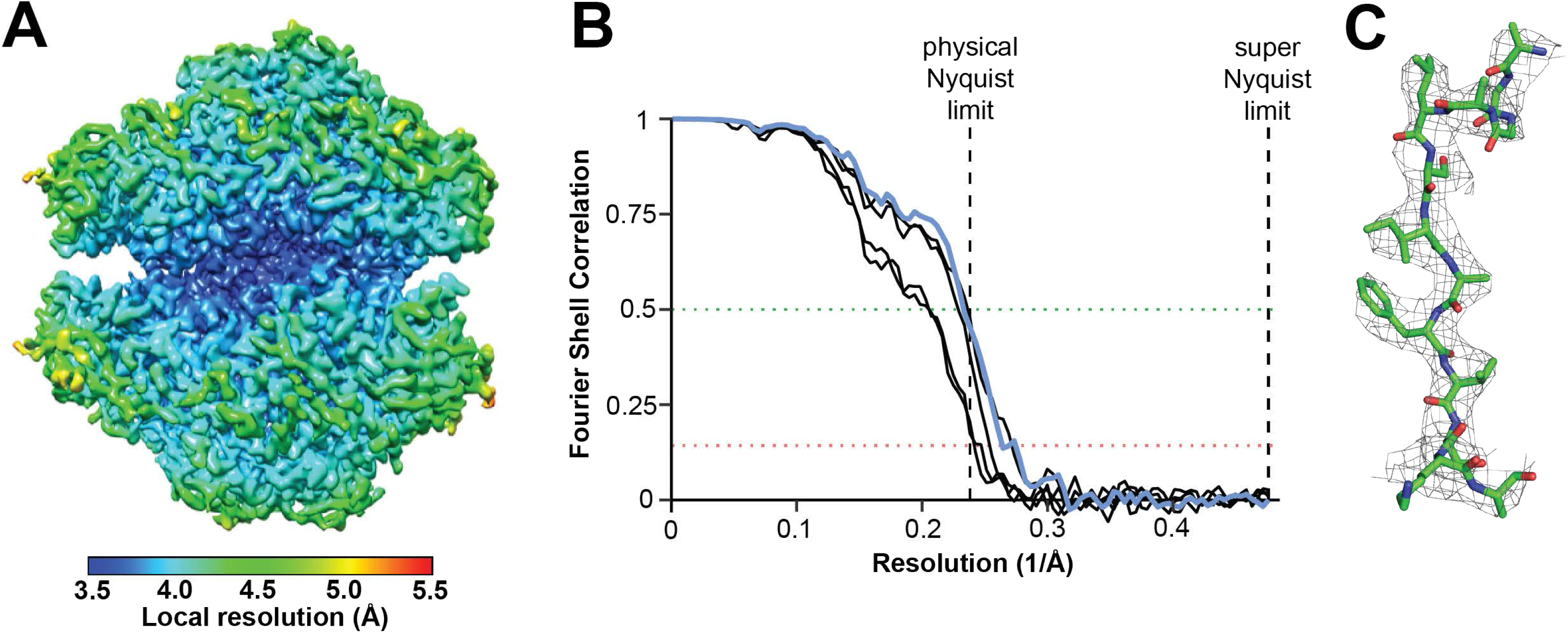
Single micrographs produce super-resolution reconstructions. A) Final 3.84 Å sharpened reconstruction generated from a subset of the particles from the micrograph shown in Figure 2A, colored by local resolution. B) FSC curves for the final particle sets used in 3D-reconstructions from five separate single-micrograph datasets. The 0.143 and 0.5 FSC cutoffs are shown as dotted lines, the physical Nyquist and super-Nyquist frequencies are shown as dashed lines. The curve for the particle set derived from the micrograph shown in Figure 2A and used for the reconstruction shown in panels *A* and *B* is shown in blue. C) Example electron density of the final sharpened 3D-reconstruction overlaid on the atomic structure. See Figure S4 for a graphical depiction of the data processing steps.

We note that the per-particle CTF refinement and Bayesian polishing algorithms implemented in RELION 3.0 (Zivanov et al., 2018) were essential for generating useful reconstructions from the single micrographs collected at 39kX magnification. Without these additional data processing steps, the resolution of the reconstructions was limited to ~8 Å (Figure S4A).

We then performed a similar analysis on the 49kX dataset, and produced a 3D-reconstruction with a 0.143 FSC resolution of 3.50 Å using 1,590 particles autopicked from a single micrograph (Figure S4B). Attempts to select a subset of these particles using 3D classification failed to improve the reconstruction.

### Comparison of data collected at higher and lower magnifications

Although lower magnification imaging results in more particle images per micrograph, these particle images have less information content in the highest resolution ranges compared to particle images acquired at higher magnification (Figure S1B). Although this consequence of DQE deterioration is well-known (McMullan et al., 2009), we sought to systematically assess the practical consequences of using different magnifications. Therefore, to compare the costs and benefits of imaging at different magnifications, we collected additional datasets using higher nominal magnifications of 63kX (1.29 Å image pixel size) and 79kX (1.035 Å image pixel size). These datasets produced 3D reconstructions with 0.143 FSC resolutions of 2.84 Å and 2.86Å (Figures S5, S6, Table 2). We note these reconstructions did not reach the physical Nyquist limit, and therefore we used particle images that were Fourier-cropped to the physical-pixel size. We selected the best single exposures from each of these higher magnification datasets and processed them to produce 3D-reconstructions as described above. The 63kX single exposure had 932 autopicked particles, which were used to generate a 3D-reconstruction with 0.143 FSC resolution of 3.85 Å (Table 2, Figure S7A). The 79kX single exposure had 670 autopicked particles, which were used to generate a 3D-reconstruction with 0.143 FSC resolution of 4.06 Å (Table 2, Figure S7B).

We considered the possibility that at higher magnifications, the lower resolutions of the reconstructions produced from single micrographs arose from an insufficient defocus range for the particles. Single-particle experiments require particles to be imaged at varying defocus distances in order to compensate for the information loss (“zeros”) at certain frequencies due to modulation of the image by the contrast transfer function (CTF) of the instrument’s electron beam. Typically, this is compensated for by acquiring exposures at different defocus distances during data collection. In the single image collected at 39kX, most of the particles appear to be positioned across a ~150 nm range of defocus distances in the image (Figure 5A), perhaps due to the local topography of the ice layer in the hole. For the single 49kX exposure used for reconstruction, the bulk of the particles spanned a defocus range of ~60 nm, and for the 63kX and 79kX single exposures most of the particles on each exposure spanned a narrower range of ~40 nm (Figure 5A). The smaller defocus ranges at higher magnifications made us concerned that the resolution of these reconstructions could be limited by CTF zero effects.

**Figure 5:**
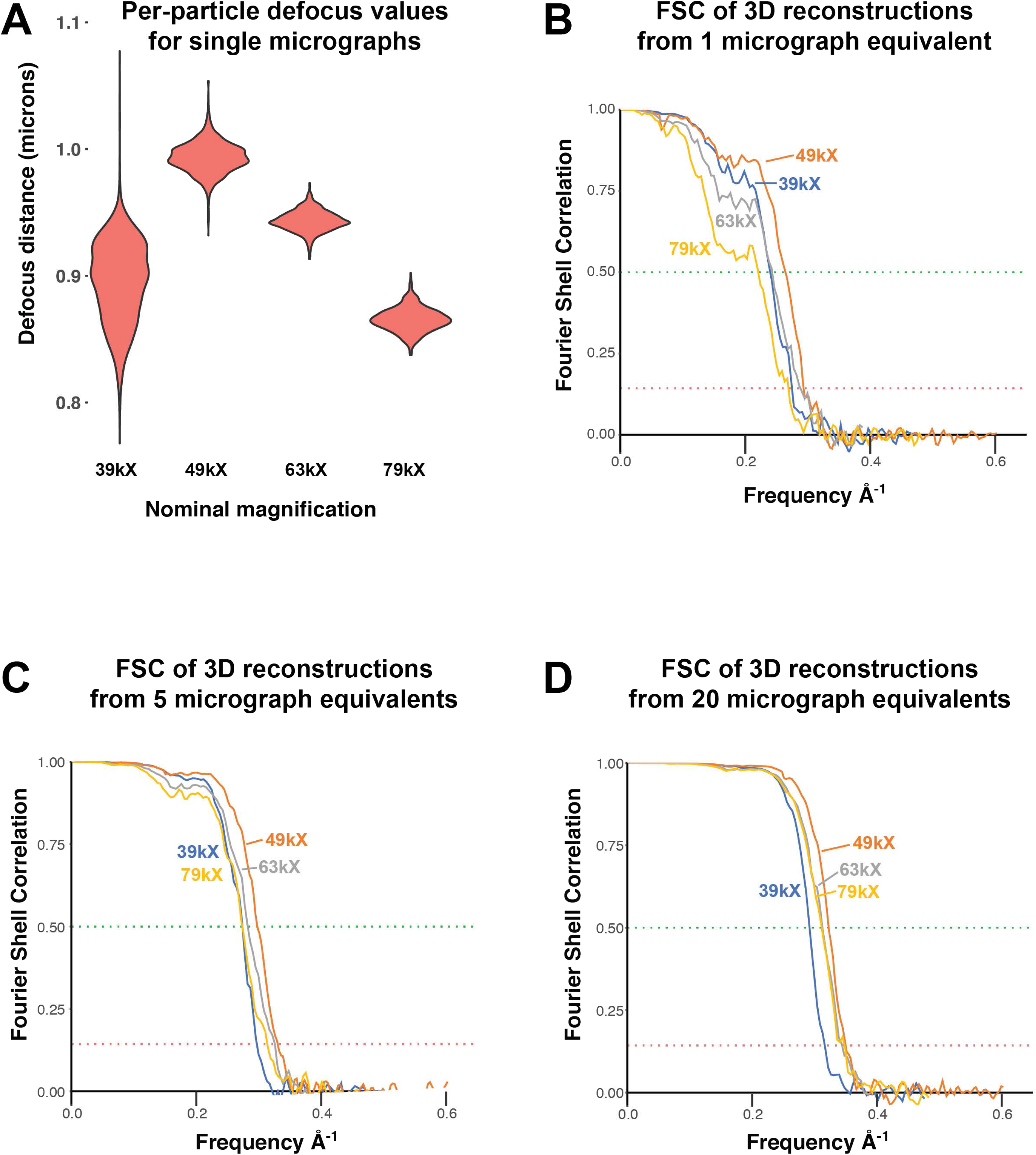
Comparing datasets collected at four different magnifications. A) Violin plots indicated the final refined defocus values of all urease particles in each 3D reconstruction generated from a single micrograph at the indicated magnification. B) FSC curves for 3D reconstructions of urease produced from the average number of particles imaged in a single micrograph at the indicated magnification. C, D) Same as (B), but for 5 and 20 micrographs, respectively.

Therefore, in order to obtain particles with a wider range of defocus while examining the effects of varying particle numbers obtained at different magnifications, we re-processed the data in a systematic way. We processed the first ~100 micrographs collected at each magnification, again using per-particle CTF refinement, Bayesian polishing, and 3D classification to obtain high quality reconstructions. We then selected random subsets of particles from each of these high-quality particle sets, choosing numbers of particles equivalent to those expected to be imaged in 1, 5, or 20 micrographs (“micrograph equivalents”) at each magnification. We then refined these particle subsets to determine the resolution of the resulting reconstructions (Figures 5B-D). This approach has the caveat that it is difficult to control for variations in sample quality among the datasets. However, this concern is mitigated by three factors: 1) the high overall quality of the sample; 2) our use of large starting numbers of particles for each magnification, which are then processed to obtain high quality sets of particles used for subsequent subset selection; 3) The 63kX and 79kX dataset were obtained from the same grid square, providing at least one very close comparison. However, we cannot completely exclude the possibility that local differences in sample quality between the datasets play a role in the observations.

Comparing the 0.5 FSC resolution of the reconstructions produced with the same number of “micrograph equivalents” at the four different magnifications, two main results emerge. First, as expected, the best resolution attainable at the lowest magnification is clearly worse than that of the three higher magnifications (Figure 5D, 6A). The straightforward explanation of these results is that at the lowest magnification (39kX), with a physical pixel size of 2.1 Å, the DQE at frequencies above ~3.1 Å is low enough to prevent the resolution of the reconstruction from surpassing this value.

Correspondingly, the fact that reconstructions produced from the three highest magnifications plateau at ~2.8 Å indicates that DQE is unlikely to be significantly limiting the useable information content for these three magnifications. Therefore, we suspect other factors, such as the imaging optics and the sample itself likely limit the resolution reached by these reconstructions.

This interpretation is further supported by plotting the same data differently, in which the 0.5 FSC resolutions of the reconstructions are compared by particle number for each magnification (Figure 6B). This analysis shows that for the three highest magnifications, the curves of particle number vs. resolution are roughly equivalent, although as expected from DQE limitations there is a slight trend toward higher resolution reconstructions at higher magnifications. In contrast, for the lowest magnification the resolution vs. particle number curve is significantly lower. This is the expected result if low DQE at high resolutions is limiting, which appears to be the case at 39kX magnification. At the three highest magnifications, the amount of high-resolution information present in the particle images appears to be quite similar, and therefore at 49kX the quality of the reconstruction does not appear to be limited by the detector DQE.

**Figure 6:**
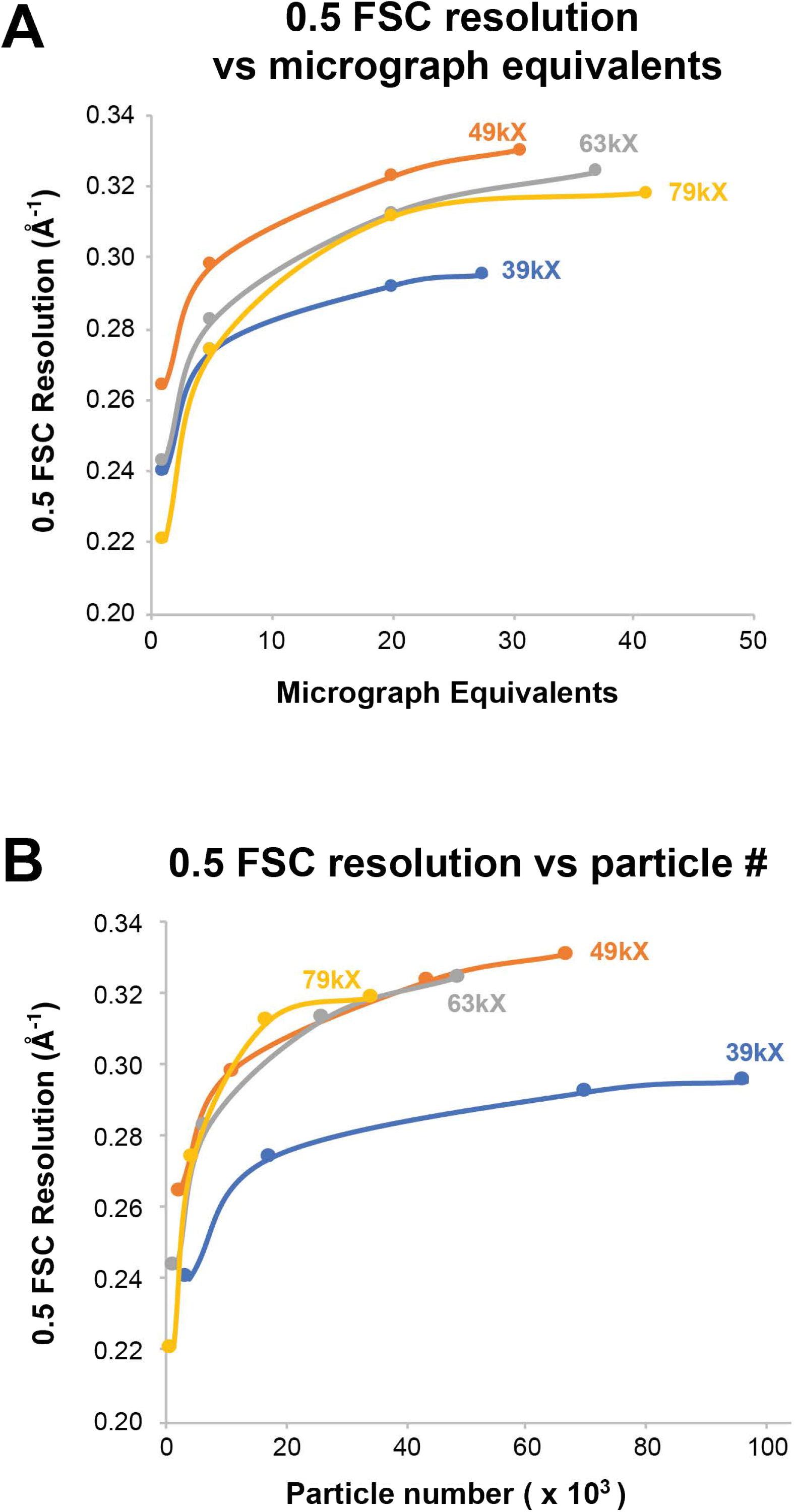
Lower magnification imaging can produce equivalent reconstructions with fewer micrographs. A) Plot of 0.5 FSC resolution values versus micrograph equivalents (average number of particles imaged per micrograph) for urease 3D reconstructions produced from grids imaged at the indicated magnifications. B) Plot of 0.5 FSC resolution values versus particle number for urease 3D reconstructions produced from grids imaged at the indicated magnifications.

Examining the FSC curves of the reconstructions produced from low (1 and 5) micrograph equivalents at the three highest magnifications (Figures 5B,C, 6A) reveals the second major result: for these small datasets, there was an observable benefit obtained by using lower magnifications. The FSC curve for the reconstruction from the 1 and 5 micrographs 49kX datasets (1.66 Å physical pixel) were better than those of the 63kX datasets (1.24 Å physical pixel), which were better than those of the 79kX datasets (1.035 Å physical pixel). The explanation for this observation arises from the interpretation, described above, that the quality of particle images obtained was very similar between these three magnifications, and the key constraint of these reconstructions was the limiting numbers of particles. The higher number of particles obtained at the lower magnifications provided more averaging power and higher resolution reconstructions.

Taken together, our results support the idea that the loss in high-resolution signal that accompanies a reduction in magnification because of the detector DQE does not always have a practical impact on the resulting reconstructions. Furthermore, under certain circumstances, there appears to be a benefit to using a lower magnification, meaning that fewer exposures are needed to generate an equivalent reconstruction.

## Discussion

In spite of the fact that super-resolution cryo-EM imaging has been available since the introduction of the K2 detector in 2012, no super-resolution single particle reconstructions have been reported in the literature. We now demonstrate that super-resolution reconstructions can be produced with 0.143-FSC resolutions approaching 1.4x the physical Nyquist frequency. We found the super-resolution information significantly improved reconstructions that were otherwise limited by the physical Nyquist limit.

We were surprised at how easily the physical Nyquist limit was surpassed, but it is important to note that jack bean urease represents an ideal sample that exhibits low flexibility and high symmetry.

Because the DQE of detectors decreases at higher frequencies (McMullan et al., 2014, 2009; Milazzo et al., 2010; Ruskin et al., 2013), high magnifications are generally required to obtain robust signal in the high-resolution range. Our analysis highlights the fact that, as is already known, there are multiple factors that limit the resolution of reconstructions, and the DQE of the detector is only one of them (Ruskin et al., 2013). To illustrate this idea, consider a contrived case in which the sample itself cannot yield information at a resolution better than 3.0 Å, perhaps because of poor image contrast arising from thick ice, or extreme and continuous structural heterogeneity. In such a case, the detector DQE at resolutions better than 3.0 Å is inconsequential. Therefore, the best magnification would be one that maximizes field of view while maintaining a reasonable DQE at 3.0 Å, rather than a higher magnification which would retain a reasonable DQE at even higher spatial frequencies that would be unlikely to contribute useful information to the reconstruction.

We found that lower magnification imaging can produce 3D-reconstructions of similar or better quality to those produced from higher magnification imaging with the same number of exposures. We explored the tradeoffs between the diminishing high-resolution signal and increasing particle numbers obtained with lower magnification imaging. We produced 3D-reconstructions from the 39kX, 49kX, 63kX, and 79kX datasets using particle numbers that would be obtained from different numbers of exposures at each magnification. This analysis directly compared 3D-reconstructions produced from the amount of data that can be collected in equal time periods using a 2-condenser lens instrument in which only a single exposure can be taken in each foil hole. We found that on a per-micrograph basis, nominal magnifications of 49kX (1.66 Å physical pixel) and 63kX (1.24 Å physical pixel) provided roughly equivalent reconstructions, whereas 39kX (2.10 Å physical pixel) and 79kX (1.035 Å physical pixel) were worse, but for different reasons. For the 39kX dataset, our results indicate that the low DQE at higher resolutions limited the resolution obtainable. For the 79kX dataset, the reduced number of particle images obtained using high magnification appeared to limit the resolution obtainable when micrographs were limiting.

An increasingly common approach that facilitates collection of large amounts of data is the use of beam-image shift (a.k.a. beam tilt or beam shift) acquisition to acquire multiple exposures per sample stage movement (Cheng et al., 2018). This is beneficial because significant time is spent waiting for stage drift to diminish after each stage movement required to center on a new hole for image acquisition. There are two approaches to beam-image shift that have been implemented, one is to take multiple exposures (as many as 10) per hole on larger (~2 μm) holes (de la Peña et al., 2018). This is only possible on 3-condenser instruments such as the Titan Krios, in which the third lens enables the formation of a very small parallel beam. With a typical 2-condenser instrument like the Talos Arctica equipped with standard apertures, the beam cannot be made small enough while maintaining parallel illumination to enable multiple exposures per hole. The other beam-image shift approach is to acquire single exposures from several holes per stage movement (Cheng et al., 2018). This is possible on both 2-condenser and 3-condenser instruments. The image distortion effects of beam-image shift can be corrected for during imaging (Glaeser et al., 2011) or data processing (Zivanov et al., 2020, 2018).

If users have access to a 3-condenser microscope, obtaining multiple high magnification exposures per hole is likely a better approach than lower magnification imaging. The potential benefits of the lower magnification approach arise for users of a 2-condenser microscope when conventional data collection strategies would not yield enough particles in the available time to generate interpretable reconstructions. When taking single exposures per hole, lower magnification captures more particle images from each hole that is imaged, so more particle images will be obtained from each hole with good ice.

Because lower magnification imaging provides more particles per image, it has potential benefits in any case where particle numbers are limiting. We have found lower magnification imaging to be beneficial for quickly screening new samples, as only a few images are needed to obtain good 2D classes and 3D-reconstructions for samples with dense particle distributions. In addition, some samples exhibit asymmetric particle distribution in holes (i.e., particles are excluded from the center of holes); in such cases a larger field of view is helpful in capturing particle images more efficiently. Furthermore, using lower magnification imaging to obtain millions of particle images may facilitate structure determination for samples with significant conformational or biochemical heterogeneity. At the least, we hope that researchers will carefully consider whether they really need to image their sample at a physical pixel size of 1.0 Å and below if the goal is a ~2.8 Å map and factors other than DQE are likely to limit the achievable resolution.

Using lower magnifications certainly has drawbacks. Lower magnifications require longer exposure times in order to maintain the optimum dose/pixel rate on the camera. This is the case for the K2 camera. However, using the K3 camera each movie exposure at 39kX only took a total of 5 seconds, which is much shorter than the amount of time (typically 30 seconds to 1 minute) spent moving between holes and waiting for stage drift to settle.

Lower magnification super-resolution imaging also results in increased data storage and data processing requirements associated with super-resolution micrographs and larger particle numbers. However, the cost of data storage continues to decline, with 8 TB hard drives currently costing only $130 USD. Our super-resolution movie exposures from the K3 are typically ~1.7 GB each if saved as gain-corrected images (and significantly smaller if not gain-corrected), so a typical super-resolution dataset will tend to be less than 2 TB in size and the cost to store each dataset is therefore < $35. Considering all the other costs associated with a cryo-EM experiment, such storage costs are relatively insignificant, and new file formats may result in either smaller image files or else the ability to capture even more information from super-resolution imaging (Guo et al., 2020). As instrument time often costs $100-$200/hour in user fees, there are strong incentives to collect as much data as possible per unit time. On the other hand, the increased processing time required to analyze more particles and larger box sizes during classification and refinement can also be significant.

The best resolution achievable for a single-particle reconstruction of a macromolecule depends upon many factors, including the properties of the molecule itself, sample preparation quality, and the microscope imaging conditions. Our results demonstrate that super-resolution 3D-reconstructions can be relatively straightforward to produce and suggest that lower magnification imaging is a viable option for data collection using 2-condenser microscopes.

## Methods

### Sample Preparation

Powdered urease from jack bean (*Canavalia ensiformis*) (Sigma-Aldrich #U0251) was solubilized in phosphate buffered saline (137 mM NaCl, 2.7 mM KCl, 8 mM Na2HPO4, 2 mM KH2PO4) at a concentration of 0.8 mg/ml, flash frozen, and stored at −80°C. Aliquots were centrifuged at 14,000g for 10 min upon thawing to remove aggregates. 3 μl of the protein solution was applied to Quantifoil R1.2/1.3 300-mesh gold-support grids that had been glow-discharged for 80 seconds at 30 mA in a Pelco EasiGlo instrument. Grids were blotted for 2.5 seconds at 4°C and 100% humidity and immediately plunge frozen in liquid ethane using a FEI Mark IV Vitrobot. Grids for both datasets were prepared at the same time under the same conditions.

### Imaging conditions

Cryo-EM data collection was performed using a Thermo Fisher Scientific Talos Arctica operated at 200 keV equipped with a Gatan K3 detector operated in counting mode with 0.5X-binning (super-resolution mode) and a Gatan BioQuantum energy filter with a slit width of 20 eV.

Microscope alignments were performed on a gold diffraction cross-grating following published procedures for a 2-condenser instrument (Herzik et al., 2017, 2019). Parallel conditions were determined in diffraction mode at the imaging magnification. Beam tilt, rotation center, objective astigmatism, and coma-free alignments were performed iteratively at the imaging magnification. A C2 aperture size was chosen so that the beam was larger than the imaging area (50 μm for 49kX and 70 μm for 39kX). The spot size was set so that the dose rate in vacuum at the detector was ~40 e^−^/physical pixel/sec. A 100 μm objective aperture was used during data collection.

SerialEM (Mastronarde, 2005; Schorb et al., 2019) was used for automated data collection of fractionated exposures (movies). The record settings were set to capture a total dose of ~24 e^−^/Å ^2^ for the 49kX dataset or ~45 e^−^/Å^2^ for the 39kX dataset, and fractionated so that the dose per frame was ~0.8 – 0.9 e^−^/Å^2^. Both the 63kX and 79kX datasets were collected using similar conditions with 50 frame movies fractionated so that the total dose was ~50 e^−^/Å^2^ and the dose per frame was ~1.0 e^−^/Å^2^. Dose rates on the detector over vacuum were ~25-30 e-/physical pixel/second.

Data for the 63kX and 79kX magnification reconstructions was collected from the same grid with over 80% of the images taken from the same square. Microscope alignments were performed at the lower magnification on a gold cross grating grid and verified on amorphous carbon after changing magnification.

### Data Processing

All cryo-EM data processing was performed within RELION 3.0 (Zivanov et al., 2018), and software was maintained by SBGrid (Morin et al., 2013). All 3D refinements and 3D-classifications imposed D3 symmetry (six asymmetric units per molecule).

Chimera (Pettersen et al., 2004) was used for visualization of reconstructions and determination of the threshold values used to create masks. Masks were created as follows: mask thresholds were determined by examination of unfiltered half-maps low-pass filtered to 15 Å, identifying the lowest threshold that removed most “dust” (solvent noise); masks were then created in RELION by low-pass filtering the reconstruction to 15 Å, applying the threshold, then adding a 6 pixel soft edge.

#### 49kX magnification (physical pixel size of 1.66 Å)

Frames from 240 movies were aligned and dose-weighted with MotionCor2 (Zheng et al., 2017) using 30 (6×5) patches, the default B-factor of 150, and no binning. Defocus values were calculated for the non-dose-weighted micrographs with GCTF (Zhang, 2016). 147 particles were manually picked from 6 micrographs and 2D classified to generate 4 templates for autopicking. 11,000 particles were autopicked from 16 micrographs and subjected to 3D-refinement to produce an initial 5.7 Å reconstruction, using the crystal structure of urease, PDB: 3LA4 (Balasubramanian and Ponnuraj, 2010), lowpass-filtered to 60 Å as a reference model. This reconstruction was then used as a 3D template for autopicking the entire dataset. 358,703 particles from all 240 dose-weighted, motion-corrected micrographs were extracted and binned 2x to the physical pixel size of 1.66 Å, then subjected to 3D refinement resulting in a reconstruction with a masked 0.143 FSC resolution of 3.32 Å (see Figure S2 for FSC curve). The aligned particle images were then re-extracted using the unbinned super-resolution pixel size of 0.83 Å. 3D-refinement of this unbinned particle set produced a reconstruction with a masked 0.143 FSC resolution of 3.1 Å, surpassing the physical pixel Nyquist limit of 3.32 Å. The data was then CTF refined (per-particle defocus estimation), Bayesian polished, and subjected to 3D-classification (k=3) keeping alignments fixed. The best class contained 128,947 particles and after 3D refinement produced a 2.8Å resolution reconstruction. A second round of CTF refinement, 3D-classification, and 3D-refinement resulted in a final reconstruction containing 56,038 particles with a masked 0.143 FSC resolution of 2.77 Å. See Figure S2 for a graphical flowchart of the data processing steps.

#### 49kX Single exposure

1,590 particles picked with a LoG blob were extracted from a single motion corrected micrograph with a super resolution pixel size of 0.83Å/pixel and refined using the map from the full dataset lowpass filtered to 60Å as a reference model with a 200Å spherical mask and D3 symmetry. The resulting map had an FSC estimated resolution of 4.03Å. Two rounds of per particle CTF correction and re-refinement increased the estimated resolution to 3.84Å. A final refinement after Bayesian polishing further improved the resolution estimate to 3.50Å. Attempts to improve the map using 3D classification were unsuccessful.

#### 39kX magnification (physical pixel size of 2.10 Å)

Frames from 284 movies were aligned and dose-weighted with MotionCor2 (Zheng et al., 2017) using the unbinned super-resolution pixel size and 30 (6×5) patches. Defocus values were estimated by CTF-fitting the non-dose-weighted micrographs with GCTF (Zhang, 2016) using information up to a resolution limit of 12 Å (using the argument “--resH 12”). 21 micrographs with GCTF maximum resolution estimates worse than 6 Å were discarded. 972,576 particles were picked from the resulting 263 micrographs using LoG autopicking in Relion. The selected particle images were extracted with a binned pixel size of 2.58 Å and 2D-classified to remove junk particles. All 2D classes that appeared to contain at least some good particles were selected resulting in 845,377 particles in 21 classes. The selected particles were refined to a generate a 3D reconstruction with resolution of 5.16 Å. Per-particle CTF refinement was then performed and the particle images were re-extracted with an unbinned super-resolution pixel size of 1.05 Å. 3D-classification with alignments was performed to reduce the number of particles in order to speed data processing. The resulting classes were essentially indistinguishable. A class of 299,603 particles was subjected to iterative rounds of 3D refinement and CTF refinement until the masked 0.143 FSC resolution converged at 3.19 Å. The output was then 3D-classified without alignments producing a class containing 111,111 particles that after an additional round of 3D refinement yielded no further improvement. Subsequent Bayesian particle polishing and 3D refinement resulted in a final reconstruction with a masked 0.143 FSC resolution of 3.06 Å. Further attempts at 3D classification and CTF refinement did not improve the resolution. See Figure S3 for a graphical flowchart of the data processing steps.

#### 20 micrograph 39kX magnification dataset

78,970 unbinned autopicked particles were extracted from the first 20 micrographs of the 39kX magnification dataset. Using an approach similar to that described above, 41,938 particles were used to generate a 3D reconstruction with a final masked 0.143 FSC resolution of 3.34 Å.

#### 5 micrograph 39kX magnification dataset

19,964 unbinned autopicked particles were extracted from the first 5 micrographs of the 39kX magnification dataset. Using an approach similar to that described above, 12,582 particles were used to generate a 3D reconstruction with a final masked 0.143 FSC resolution of 3.60 Å.

#### Single exposures from 39kX magnification dataset

Initially, the two micrographs with the lowest estimated defocus values (approximately 0.8 μm and 0.9 μm underfocus) displaying good contrast among the first 20 micrographs of the 39kX magnification dataset were selected and processed independently using an approach similar to that described above. From the 0.9 μm underfocus micrograph, 3,912 particles were autopicked and processed to generate a 3D-reconstruction with a masked 0.143 FSC resolution of 3.84 Å using 2,967 particles selected after 3D classification. From the 0.8 μm underfocus micrograph, 3,830 particles were autopicked and processed to generate a selected after 3D-reconstruction with a masked 0.143 FSC resolution of 4.12 Å using 3,160 particles selected after 3D classification. Subsequently, we chose three more single exposures to process in a similar manner. See Figure S4 for a graphical flowchart of the data processing steps leading to the 3.84 Å reconstruction.

#### 63kX magnification (physical pixel size of 1.29 Å)

Frames from 97 movies were aligned and dose-weighted with MotionCor2 (Zheng et al., 2017) using the unbinned super-resolution pixel size and 30 (6×5) patches. Defocus values were estimated by CTF-fitting the non-dose-weighted micrographs with GCTF (Zhang, 2016) using information up to a resolution limit of 12 Å (using the argument “--resH 12”).

85,603 particles were picked from the micrographs using LoG autopicking in Relion. The selected particle images were extracted with a Fourier-binned pixel size of 1.29 Å and refined to a generate a 3D reconstruction with resolution of 3.60 Å. Two rounds of per-particle CTF refinement followed by Bayesian polishing and subsequent CTF refinement was then performed resulting in a map with an FSC estimated resolution of 2.84Å. 3D-classification without alignments was performed with 3 classes and T=10 to select a class containing 56.6% or 48,716 particles. A subsequent refinement of the resulting particles produced a map with the same masked 0.143 FSC resolution of 2.84 Å. Further classification, refinement and CTF refinement did not improve the reconstruction. See Figure S5 for a graphical flowchart of the data processing steps.

#### 79kX magnification (physical pixel size of 1.035 Å)

Frames from 100 movies were aligned and dose-weighted with MotionCor2 (Zheng et al., 2017) using the unbinned super-resolution pixel size and 30 (6×5) patches. Defocus values were estimated by CTF-fitting the non-dose-weighted micrographs with GCTF (Zhang, 2016) using information up to a resolution limit of 12 Å (using the argument “--resH 12”).

68,566 particles were picked from the micrographs using LoG autopicking in Relion. The selected particle images were extracted with a Fourier-binned pixel size of 1.035 Å and refined to a generate a 3D reconstruction with resolution of 3.41 Å. Two rounds of per-particle CTF refinement followed by Bayesian polishing and subsequent CTF refinement was then performed resulting in a map with an FSC estimated resolution of 2.87Å. 3D-classification without alignments was performed with 3 classes and T=10 to select a class containing 50.4% or 34,605 particles. A subsequent refinement of the resulting particles produced a map with the an equivalent 0.143 FSC resolution of 2.86 Å. Further classification, refinement and CTF refinement did not improve the reconstruction. See Figure S6 for a graphical flowchart of the data processing steps.

### Direct comparison of 39kX particle images with and without super-resolution information

The final set of ~110,000 particles used to produce the 39kX reconstruction were re-extracted and Fourier-cropped to the physical pixel size of 2.1 Å (super-resolution pixel size: 1.05 Å/pixel, extraction box-size: 224 pixels, Fourier-cropped box-size: 112 pixels). The resulting particle images were subjected to standard 3D-refinement. The refined particles were then extracted again, both with and without Fourier-cropping to the physical pixel size. Half-map reconstructions were then produced for both the cropped and super-resolution particle sets and used for automated postprocessing. Note that we did not perform Bayesian polishing on these particles in order to do a direct comparison with and without super-resolution information.

### A note about particle image box sizes used for processing

The 49kX dataset was originally processed using a box size corresponding to ~318Å per edge. Particles from the other three datasets were processed using a box size of ~230Å per edge. To test whether or not the larger box size used in the 49kX dataset contained information contributing to reconstruction resolution estimates, the particle stack from the best 49kX reconstruction were re-extracted with a ~230Å box size and subjected to 3D autorefinement following Bayesian polishing. There was no change in the estimated resolution or quality of the 3D reconstruction.

### Atomic model building and real-space refinement

The final sharpened masked electron density map was used to build an atomic model of jack bean urease by modifying the crystal structure (Balasubramanian and Ponnuraj, 2010) in Coot (Emsley and Cowtan, 2004; Emsley et al., 2010). Real-space refinement (Afonine et al., 2018b) was performed and validated in Phenix (Moriarty et al., 2009; Adams et al., 2010; Afonine et al., 2018a; Williams et al., 2018) using the same map. Hydrogen atoms were included during refinement but removed from the models prior to final validation. We used “map-to-model” (Terwilliger et al., 2018) and “chain comparison” within Phenix to determine the fraction of urease residues that could be successfully modeled automatically. Figures depicting atomic models and electron density were generated with PyMol.

### Databases

3D-reconstructions have been deposited in the EMDB and will be released upon publication: 49kX reconstruction, EMD-20016; 39kX reconstruction, EMD-20213; single-micrograph reconstruction, EMD-20214. The 2.77 Å atomic model of urease refined against the 49kX reconstruction has been deposited in the RCSB database with PDB code 7KNS and will be released upon publication. Raw movie files will be available in the EMPIAR database upon publication.

## Acknowledgements

We thank Lena Kourkoutis and Mariena Silvestry-Ramos for critical reading of the manuscript. We thank Joel Meyerson, Chris Booth, and Niko Grigorieff for very helpful discussions. This work was supported by NIH/NIGMS grants R01GM098621, R01GM116942, and R35GM136258 to J.C.F, and NIH training grant T32GM007273 and a Ford Foundation Fellowship to J.R.F. This work made use of the Cornell Center for Materials Research (CCMR) Shared Facilities which are supported through the NSF MRSEC program (DMR-1719875). We thank the Cornell University Office of the Provost for funding the acquisition and maintenance of the microscope, and CCMR staff for advice and support.

## Author Contributions

JRF: Conceptualization, Methodology, Formal Analysis, Investigation, Writing, Revising, Visualization.

KAS: Conceptualization, Methodology, Formal Analysis, Investigation, Revising.

JCF: Conceptualization, Methodology, Formal Analysis, Writing, Revising, Visualization, Supervision, Funding Acquisition.

---

**Figure S1:**
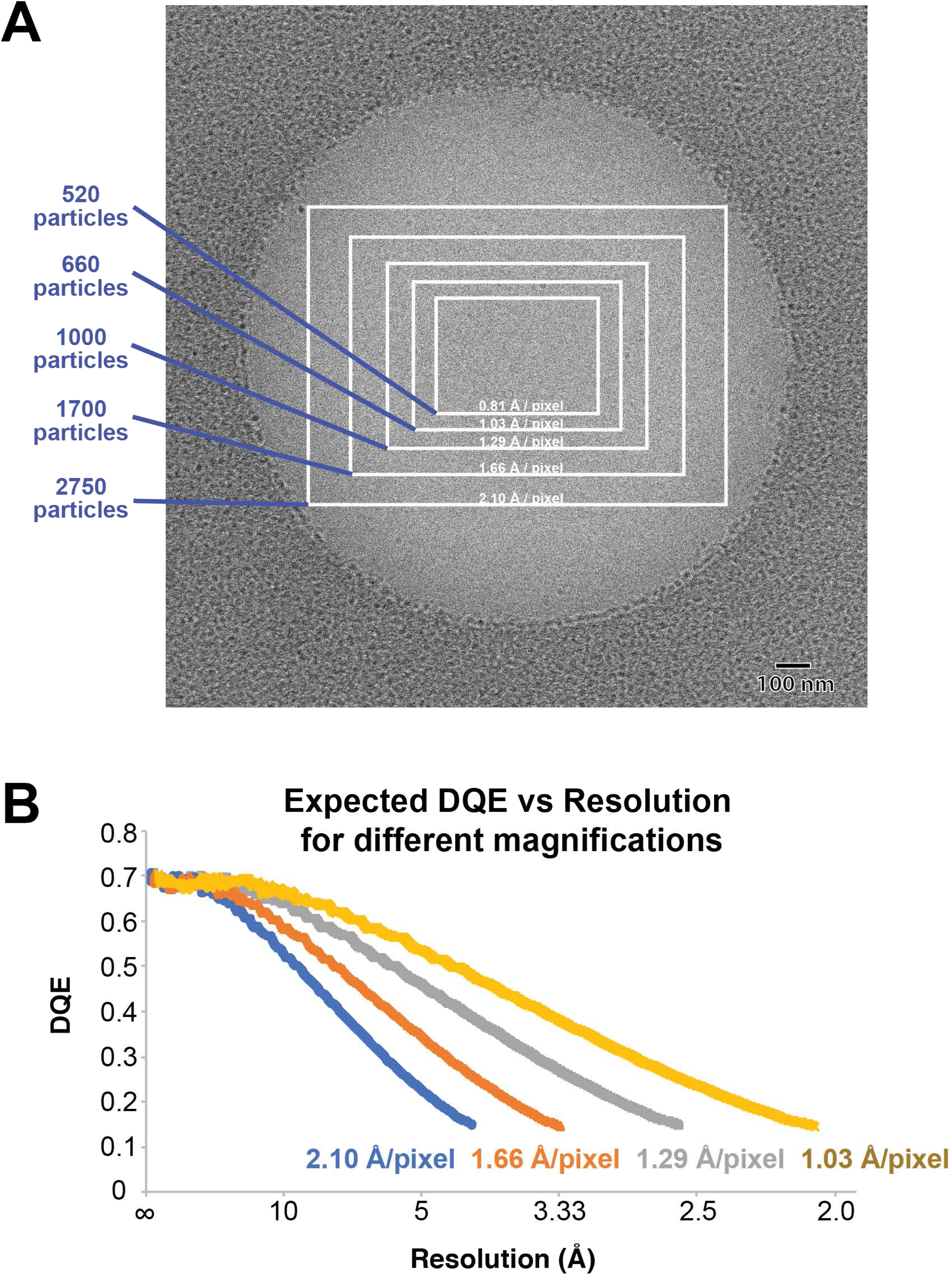
Impact of magnification on field of view. A) An image of a R1.2/1.3 Quantifoil hole with depicted imaging areas captured by the K3 detector at the indicated magnified image pixel sizes. Note, the actual hole diameter is ~1.5 μm. Relative particle numbers are shown for each field of view, assuming a uniform particle distribution (see also Table 1). The second largest field of view size was used for experiments presented in Figure 1 and the largest field of view size was used for experiments presented in Figures 2 and 3. B) Plot of DQE vs. resolution for different pixel sizes. The DQE values used were previously determined for the K2 detector (Ruskin et al., 2013) (https://grigoriefflab.umassmed.edu/dqe_data_files). These curves illustrate how the expected DQE at a given resolution (1/frequency) shifts with changes in magnification.

**Figure S2:**
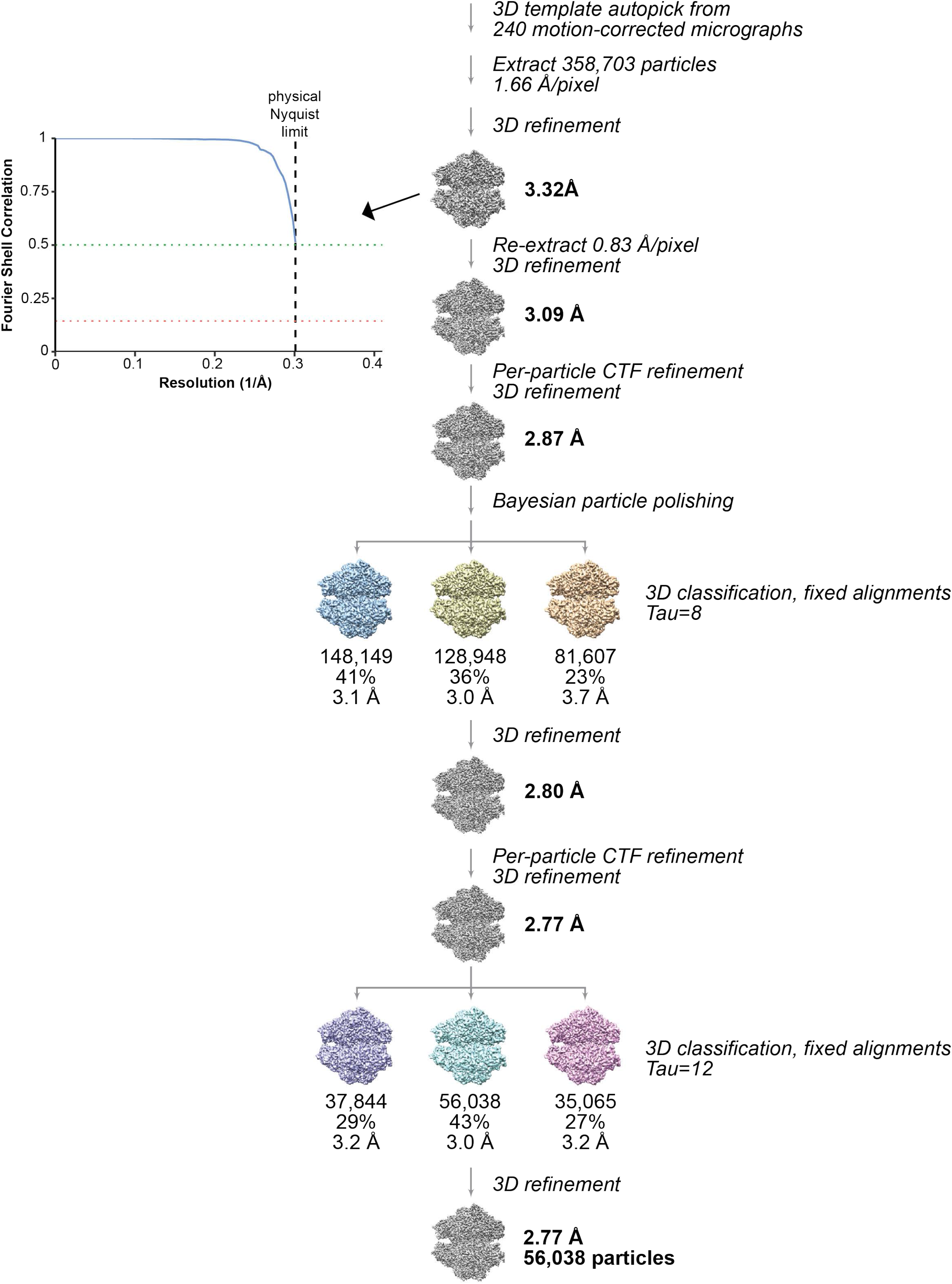
Data processing flowchart for the 49kX super-resolution dataset. See Methods for details and description.

**Figure S3:**
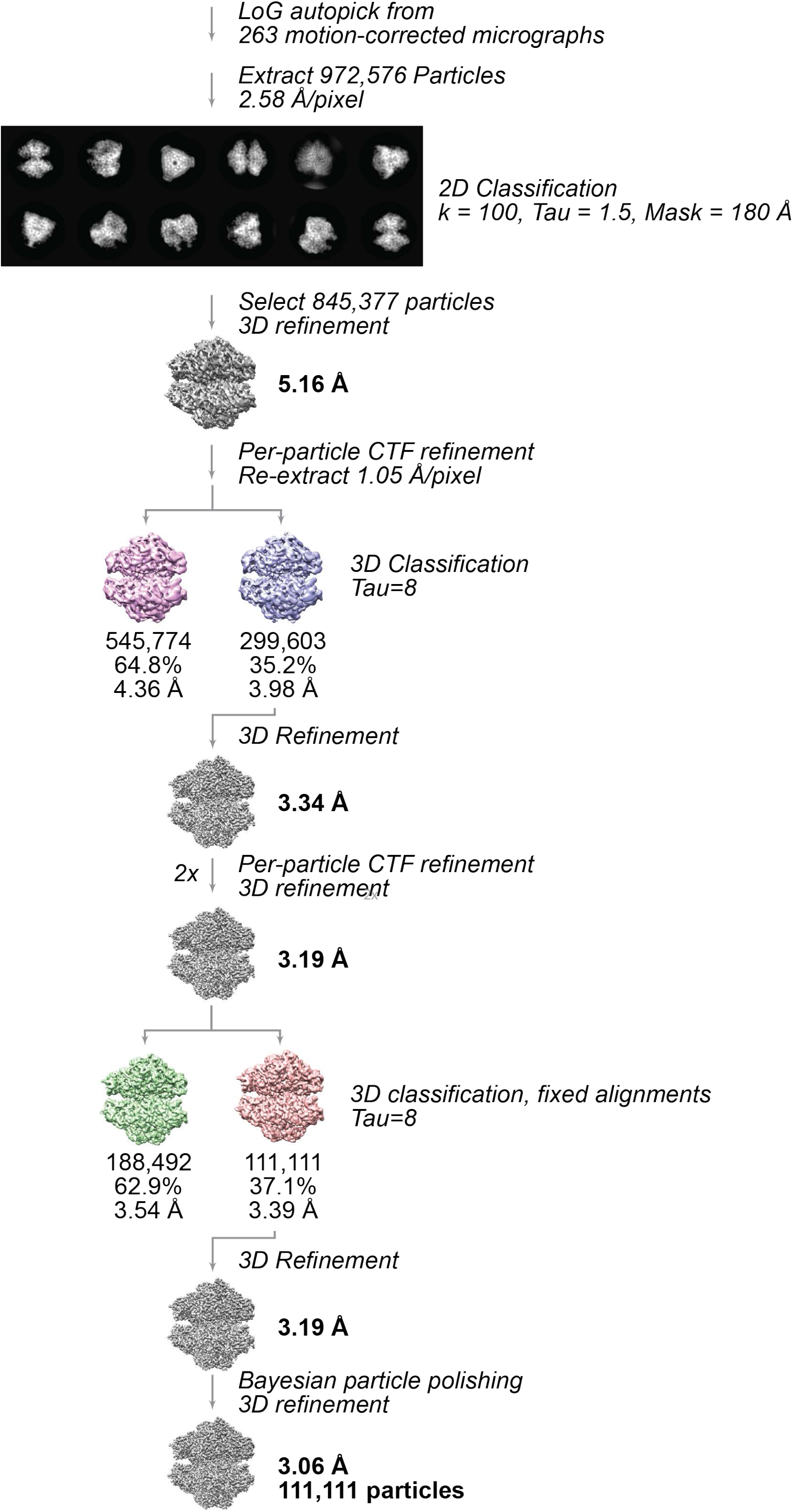
Data processing flowchart for the 39kX super-resolution dataset. See Methods for details and description.

**Figure S4:**
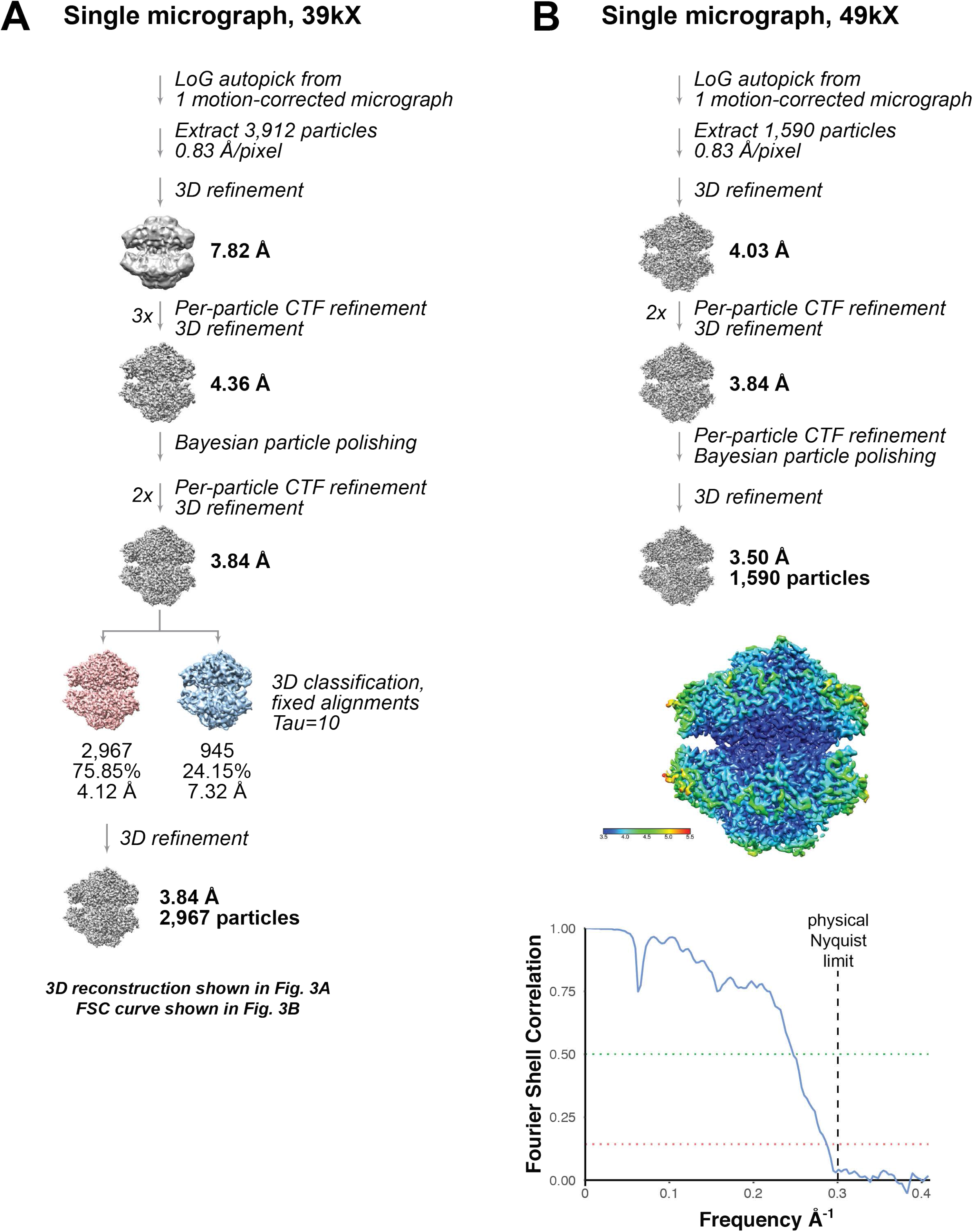
Data processing flowchart for the single micrograph super-resolution dataset. See Methods for details and description.

**Figure S5:**
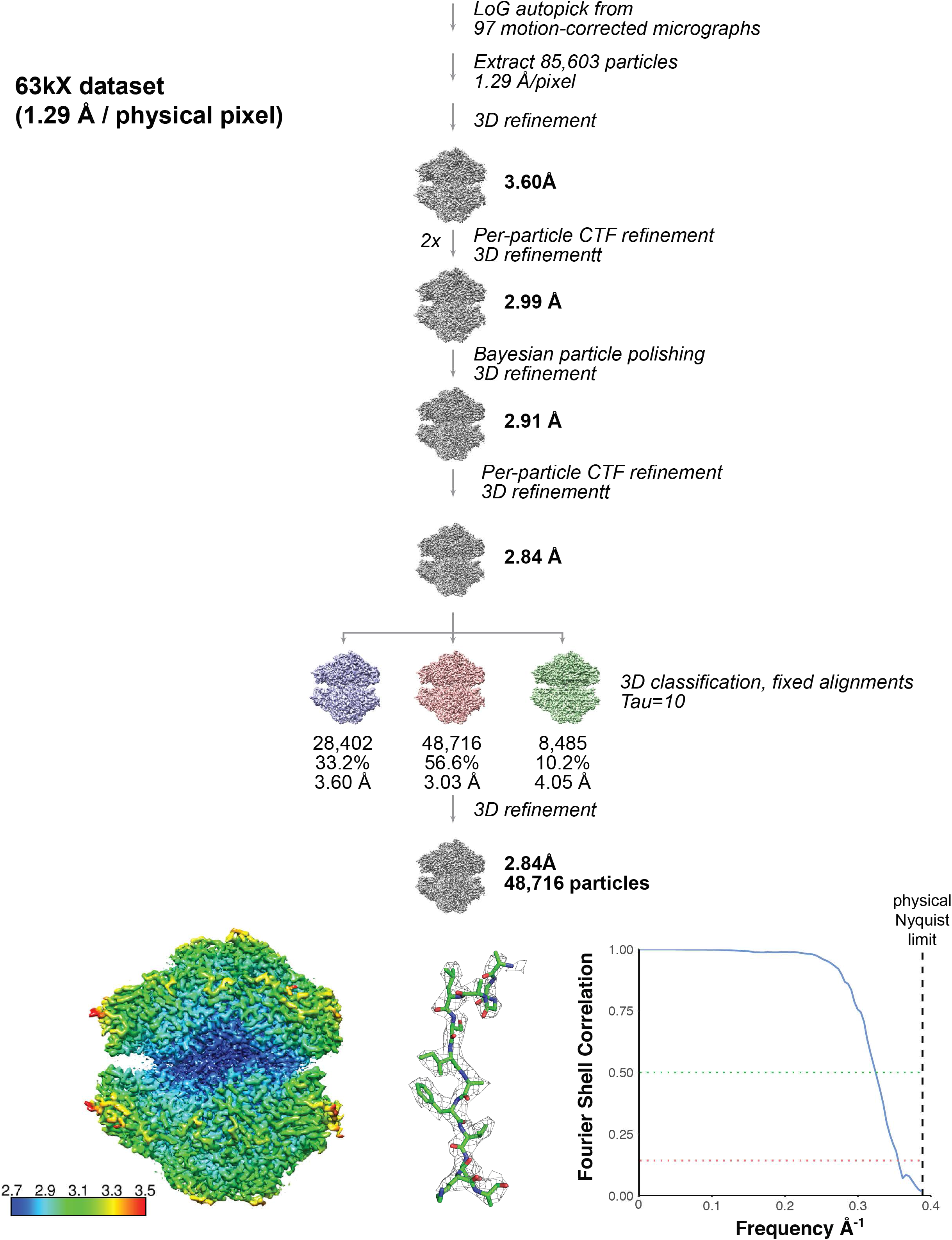
Data processing flowchart for the 63kX dataset. See Methods for details and description.

**Figure S6:**
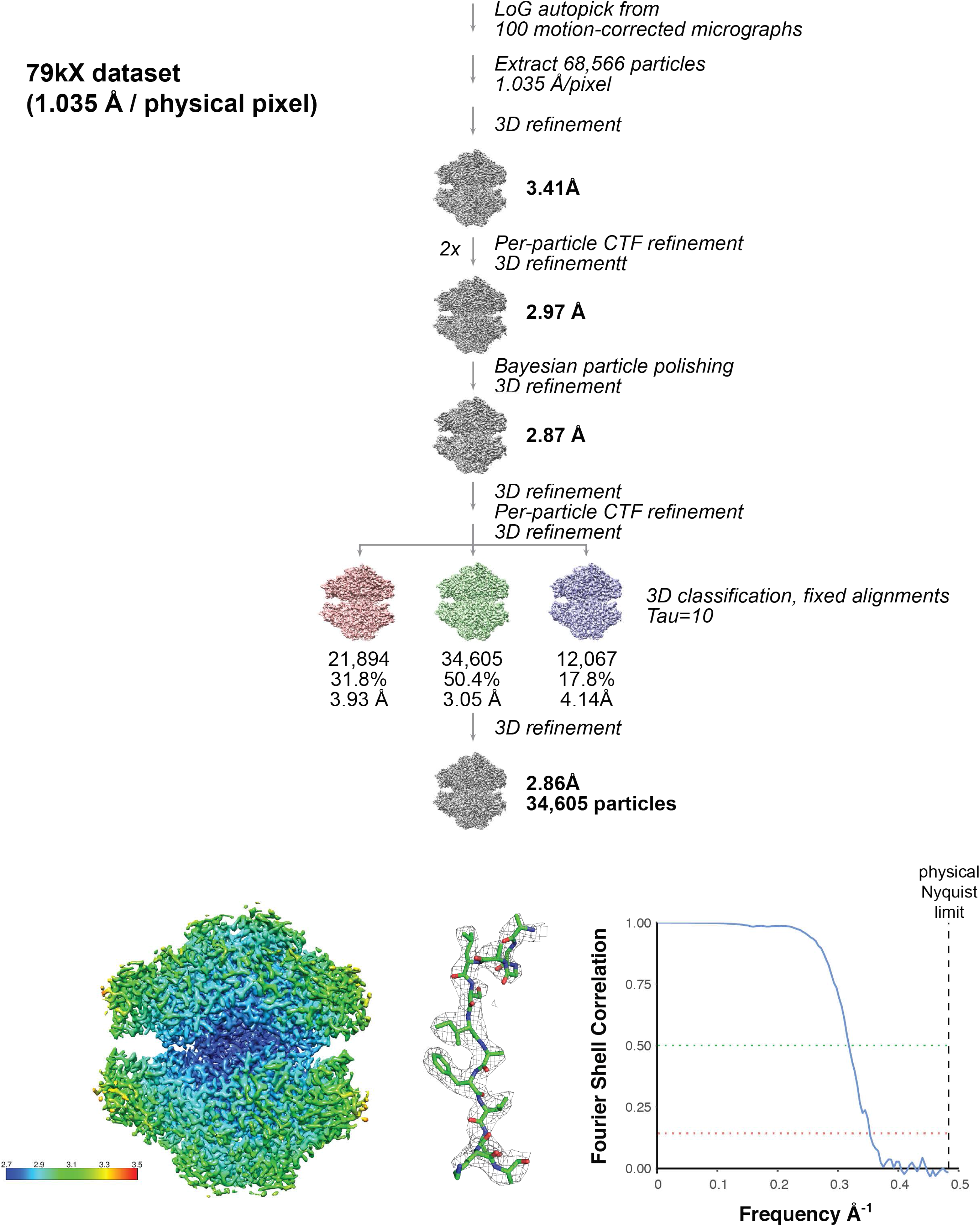
Data processing flowchart for the 79kX dataset. See Methods for details and description.

**Figure S7:**
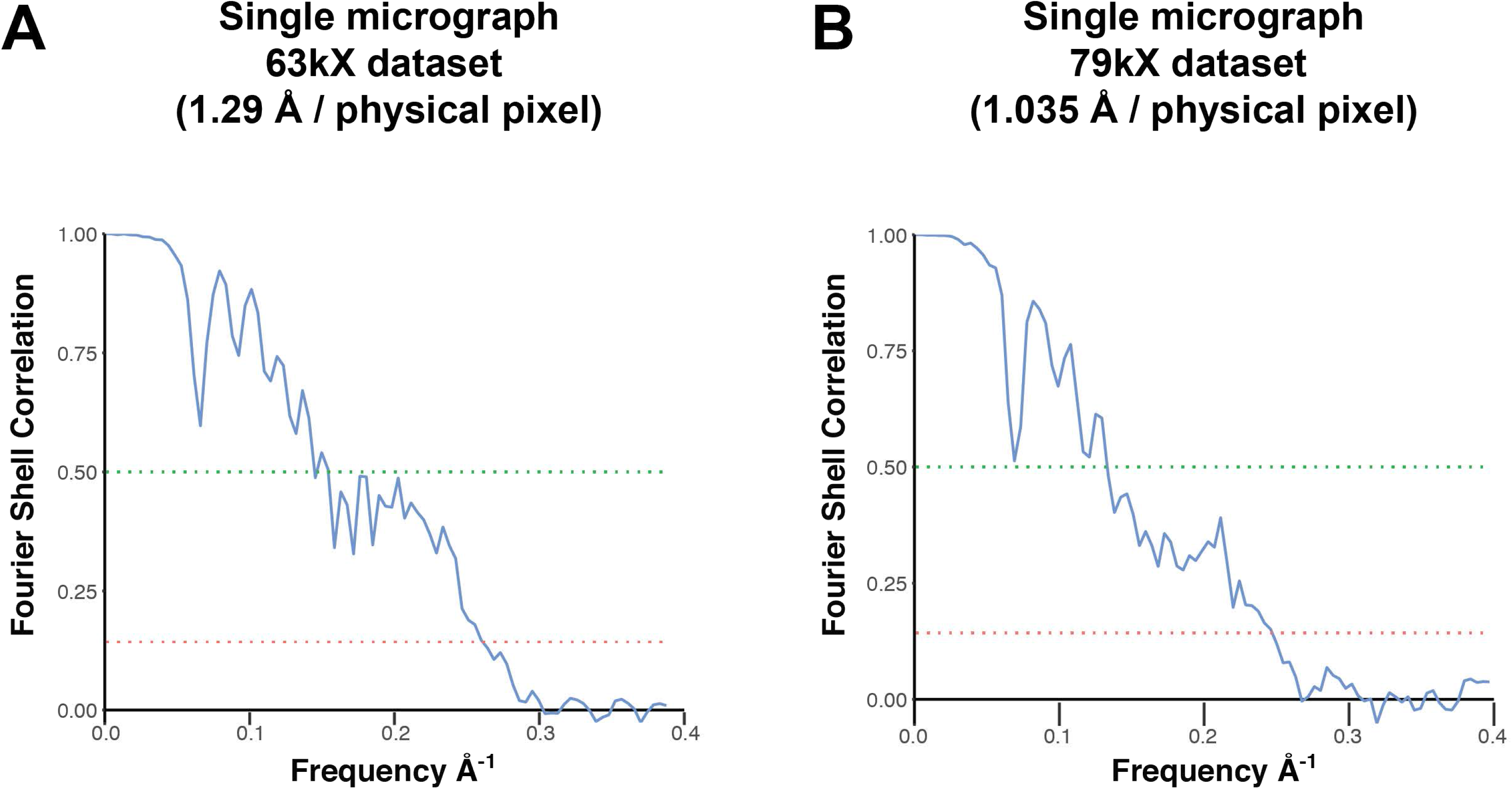
FSC curves for single micrograph 63kX and 79kX datasets. A) FSC curve for the 3D reconstruction generated from the 63kX single micrograph. B) FSC curve for the 3D reconstruction generated from the 79kX single micrograph.

